# Immuno-Scanning Electron Microscopy of Islet Primary Cilia

**DOI:** 10.1101/2024.02.16.580695

**Authors:** Sanja Sviben, Alexander J. Polino, Isabella Melena, Jing W. Hughes

**Affiliations:** Washington University Center for Cellular Imaging, Washington University School of Medicine, 660 South Euclid Ave, Saint Louis, MO, USA; Department of Cell Biology and Physiology, Washington University School of Medicine, 660 South Euclid Ave, Saint Louis, MO, USA; Department of Medicine, Washington University School of Medicine, 660 South Euclid Ave, Saint Louis, MO, USA

## Abstract

The definitive demonstration of protein localization on primary cilia has been a challenge for cilia biologists. Primary cilia are solitary thread-like projections that contain specialized protein composition, but as the ciliary structure overlays the cell membrane and other cell parts, the identity of ciliary proteins are difficult to ascertain by conventional imaging approaches like immunofluorescence microscopy. Surface scanning electron microscopy combined with immuno-labeling (immuno-SEM) bypasses some of these indeterminacies by unambiguously showing protein expression in the context of the 3D ultrastructure of the cilium. Here we apply immuno-SEM to specifically identify proteins on the primary cilia of mouse and human pancreatic islets, including post-translationally modified tubulin, intraflagellar transport (IFT) 88, the small GTPase Arl13b, as well as subunits of axonemal dynein. Key parameters in sample preparation, immuno-labeling, and imaging acquisition are discussed to facilitate similar studies by others in the cilia research community.

## INTRODUCTION

Primary cilia are surface sensory organelles that perform vital functions for the cell. The structure of the primary cilium has been studied early and often by transmission electron microscopy (TEM), which has driven most of our understanding and classification of primary versus motile cilia. However it has become clear from recent 3D ultrastructural studies of primary cilia^1–4^ that microtubule arrangements can deviate from their classic “9+0” scheme, and that functionally primary cilia are not static organelles as once thought but can rather be dynamic and exhibit motility^5–8^. Elucidating the functional significance of these cilia properties requires identification of mechanistic protein components. This has so far been done by conventionally immunofluorescence (IF) imaging which has allowed localization of axonemal dynein and motility-related proteins to human islet primary cilia^8^. However, there are pitfalls in IF colocalization studies that limit interpretation, a key one being the physical overlap among the cilium, plasma membrane, and other cell parts that confound true colocalization, and light microscopy generally lacks resolution to show sub-ciliary protein expression. As result, there has not been a good method to unambiguously identify individual proteins and their 3D distribution on primary cilia.

Scanning electron microscopy (SEM) is a high-resolution technique uniquely suited for examining the native 3D conformation of surface structures, an approach that we recently used to characterize the primary cilia axoneme of human islets^9^. In the present study, we combine our SEM protocol with antibody staining to identify specific ciliary protein components. Using 18 nm colloidal gold-conjugated secondary antibodies, we obtain clear labeling of axonemal proteins in situ and demonstrate adaptability of this technique across species in mouse and human islets.

## METHODS

### Islet preparation for SEM

Mouse islets were isolated from wildtype C57Bl/6J mice using collagenase digestion with a modified Lacy protocol. Both female and male mice were used in this study. Isolated islets were rested overnight in islet medium (RPMI 1640 with 11 mM glucose, 10% FBS, and 1% Penicillin-Streptomycin) to allow recovery of surface morphology. Human islets from a 42-year-old healthy, non-diabetic, non-obese female donor were obtained from the Integrated Islet Distribution Program (IIDP), also washed and rested overnight before use for experiments. Islets were plated on laminin-coated 12mm glass coverslips (0.5 μg laminin/cm^2^ coating overnight at 4°C, Gibco rhLaminin-521) and allowed to adhere for up to three days. For membrane extraction, adhered islets were washed 3 times in warm PBS, then incubated for 5 minutes in cytoskeleton buffer (50 mM imidazole, 50 mM KCl, 0.5 mM MgCl2, 0.1 mM EDTA, 1 mM EGTA, pH 6.8) with 0.5% Triton X-100 and 0.25% glutaraldehyde, followed by 10 minutes in cytoskeleton buffer with 2% Triton X-100 and 1% CHAPS.

### Antibody staining and immuno-gold labeling

Demembranated islets were washed in PBS 3 times, then fixed with 4% PFA in PBS for 20 minutes at room temperature with gentle shaking. Post-fixation, islets were quenched with 50 mM glycine^10^ in PBS for 15 minutes, and incubated for 30 minutes in blocking solution consisting of 1% BSA (100 mg BSA in 10 mL PBS) and 5% normal donkey serum (NDS) in PBS. Primary antibody labeling was performed at 4°C for 24 hours and then room temperature for 5 hours. Primary antibodies were used between 1:50-1:400 dilution in blocking solution, specific dilutions indicated in **Table 1**. Negative controls were performed using secondary antibody alone. After primary antibody labeling, islets were washed 5 times for 5 minutes each in blocking solution. Secondary gold labeling was performed for 1.5 hours at 1:20 dilution in blocking solution (18 nm gold donkey-anti-mouse and donkey-anti-rabbit IgG, Jackson ImmunoResearch 705-215-147, 711-215-152). Post-staining, islets were washed twice for 5 minutes each in blocking solution, then 3 times 5 minutes each in PBS, and fixed in 2% glutaraldehyde (GA) in PBS for 15 minutes.

**Table 1.** Primary antibodies for cilia immuno-SEM.

| Antibody | Antigen | Source | Host / Isotype | Dilution | Success | Figure |
| --- | --- | --- | --- | --- | --- | --- |
| AcTUB | Acetylated alpha tubulin | Proteintech 66200-1-Ig | Mouse IgG1 | 1:100 | Yes | 2-4 |
| GT335 | Polyglutamylated tubulin | Adipogen AG-20B-0020 | Mouse IgG1k | 1:400 | Yes | 5 |
| IFT88 | Intraflagellar transport 88 | NSJ Bioreagents F41236 | Rabbit IgG | 1:50 | Yes | 6 |
| Arl13b | ADP ribosylation factor-like protein 13b | Neuromab N295B/66 | Mouse IgG2a | 1:100 | Yes | 7 |
| Arl13b | ADP ribosylation factor-like protein 13b | Proteintech 17711-1-AP | Rabbit IgG | 1:100 | No, high background | 8 |
| DNAH5 | Axonemal dynein heavy chain 5 | Sigma HPA037470 | Rabbit IgG | 1:100 | Yes | 9 |
| DNAI1 | Axonemal dynein intermediate chain 1 | Neuromab 75-372 | Mouse IgG1 | 1:100 | Yes | 9 |

### Scanning electron microscopy

Coverslips containing adhered and fixed islets were stained with 0.5% osmium in PBS for 20 minutes on ice. Samples were rinsed 3 times for 10 minutes each in ultrapure water and dehydrated in a graded ethanol series (10%, 20%, 30%, 50%, 70%, 90%, 100% x3) for 5 minutes in each step. Once dehydrated, samples were loaded into a critical point drier (Leica EM CPD 300, Vienna, Austria) which was set to perform 12 CO_2_ exchanges at the slowest speed. Samples were then mounted on aluminum stubs with carbon adhesive tabs and coated with 12 nm of carbon (Leica ACE 600, Vienna, Austria). Finished samples were stored in a lab desiccator until imaging. SEM images were acquired on a Helios 5 UX DualBeam FIB-SEM platform (Fisher Scientific, Brno, Czech Republic) using SEM imaging mode at 5 kV and 0.1 nA using TLD and ICD detectors to image secondary electron and backscatter electron signal, respectively.

## RESULTS

We screened 7 primary antibodies for cilia immuno-SEM in mouse and human islets and achieved compelling labeling with 6 out of 7. These primary antibodies were used in conjunction with standard gold-conjugated secondary antibodies specific for mouse and rabbit IgG. **Table 1** lists the source and labeling conditions of all primary antibodies, typically used at 5-to 10-fold higher concentrations as for immunofluorescence imaging^8,11–13^. Some but not all primary antibodies were tested in both mouse and human islets – species indicated in figures.

Because immuno-SEM techniques had not been previously reported for primary cilia and because we anticipated difficulties in antibody labeling, we took several measures from the outset to enhance protocol performance. First, we opted to include a demembranating step in our islet sample preparation because we reasoned that membrane-stripping would expose axonemal antigens and increase the success of antibody labeling. Second, we included a gentle pre-fixation step using 0.25% glutaraldehyde (GA) prior to demembranation, which we thought would be critical for crosslinking axoneme-associated and juxta-membrane ciliary proteins such as intraflagellar transport 88 (IFT88) and the small GTPase ADP ribosylation factor-like protein 13b (Arl13b). Although a small amount of GA was expected to help preserve ultrastructural integrity, we refrained from using concentrations greater than 0.25% GA so as to protect downstream immuno-labeling. Third, we included a glycine quenching step during antibody binding to reduce sample autofluorescence and to enhance antibody-dependent signals^10^. After antibody labeling, which was done with a conventional 4% paraformaldehyde (PFA) fixation, islets undergo a final crosslink in high-concentration 2% GA before alcohol dehydration, critical point drying, and SEM imaging. We applied this workflow to both mouse and human islet samples and found it to be robust and reliable for most antibodies tested.

Multi-scale scans were acquired on intact islets adhered to glass coverslips, starting with whole-islet views at micrometer resolution to successive higher magnifications to visualize individual cilia and sub-ciliary zones at nanoscale resolution (**Figure 1**). Cilia across spatial scales were tracked by morphology and their identification aided by immunogold labeling of ciliary protein markers. Preservation of ciliary structures was excellent as most cilia axonemes remained intact, exhibiting similar lengths (4-10 µm) and diameters (200-300 nm at base) as previously characterized by us and others^9,11,14,15^. We found moderate magnification (20-35,000x) to give a good overview of the entire ciliary structure. In addition to primary cilia, other structures including microvilli and cortical cytoskeleton were also well-preserved on the islet surface.

**Figure 1.**
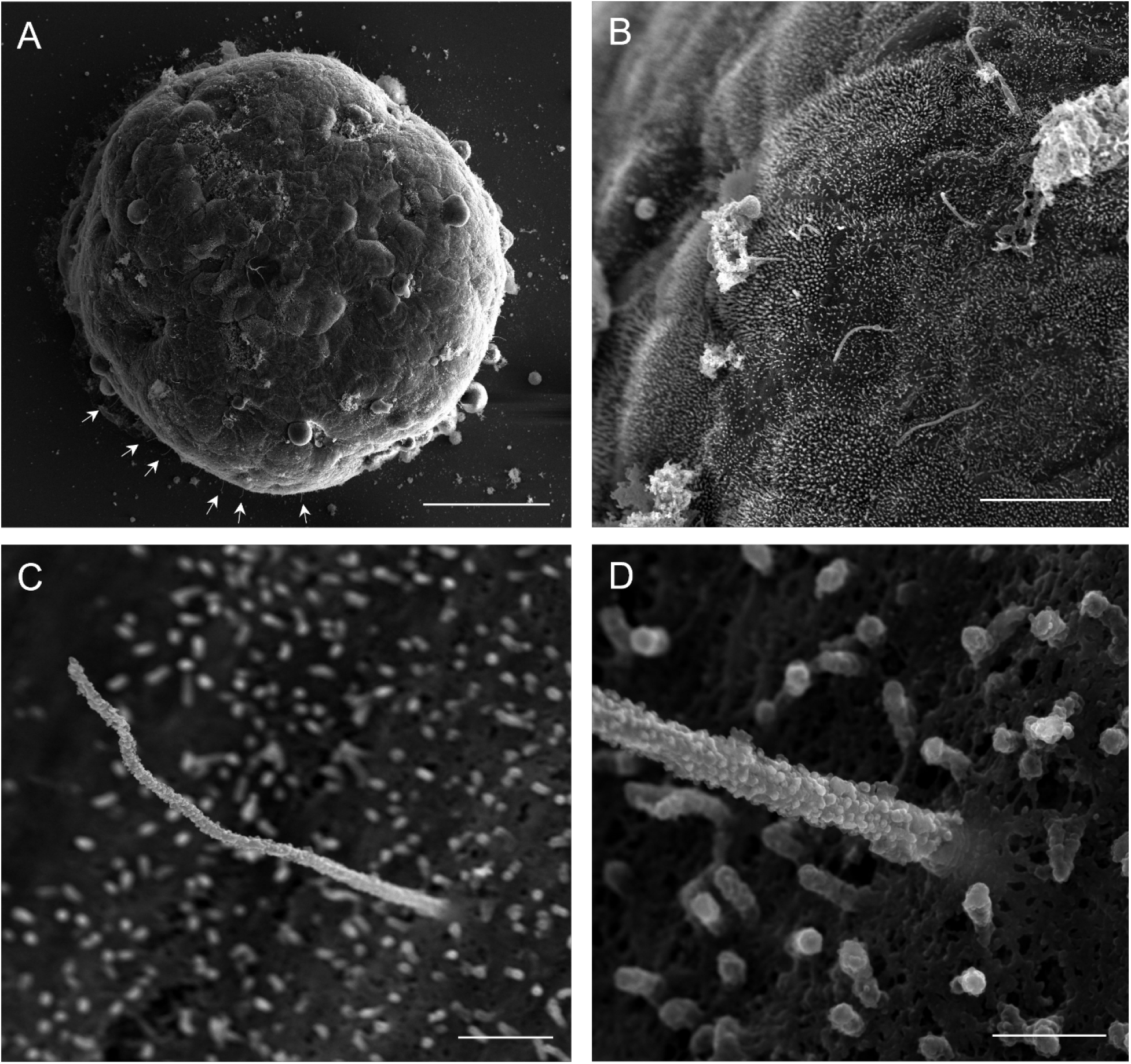
Multi-scale SEM of islet primary cilia. (**a**) An intact mouse pancreatic islet scanned at low magnification using an Everhart-Thornley detector (ETD). Small, thread-like primary cilia project from the islet surface (arrows). Scale, 50 μm. (**b**) Representative human islet surface scan at low magnification using Through Lens Detector (TLD) showing dense short microvilli, solitary primary cilia, and post-demembranation debris. Scale, 10 μm. (**c**) A human islet primary cilium at moderate magnification imaged with TLD detector. Scale, 1 μm. (**d**) High-resolution scan of the cilium base from (**c**) in high magnification imaged with TLD detector. Scale, 400 nm.

We acquired simultaneous images using multiple detectors. **Figure 2** shows images acquired with TLD detector (collects secondary electron signal for standard topography imaging), ICD detector (collects backscattered electron signal that reveals immunogold label), and Mix (overlayed TLD and ICD images). Gold particles are seen as white dots in the ICD image and overlap with cilia shape outline, confirming that imaging conditions were good and that labeling is detected. Negative control was tested for both anti-mouse and anti-rabbit secondary antibodies, which produced no labeling (**Figure 3**).

**Figure 2.**
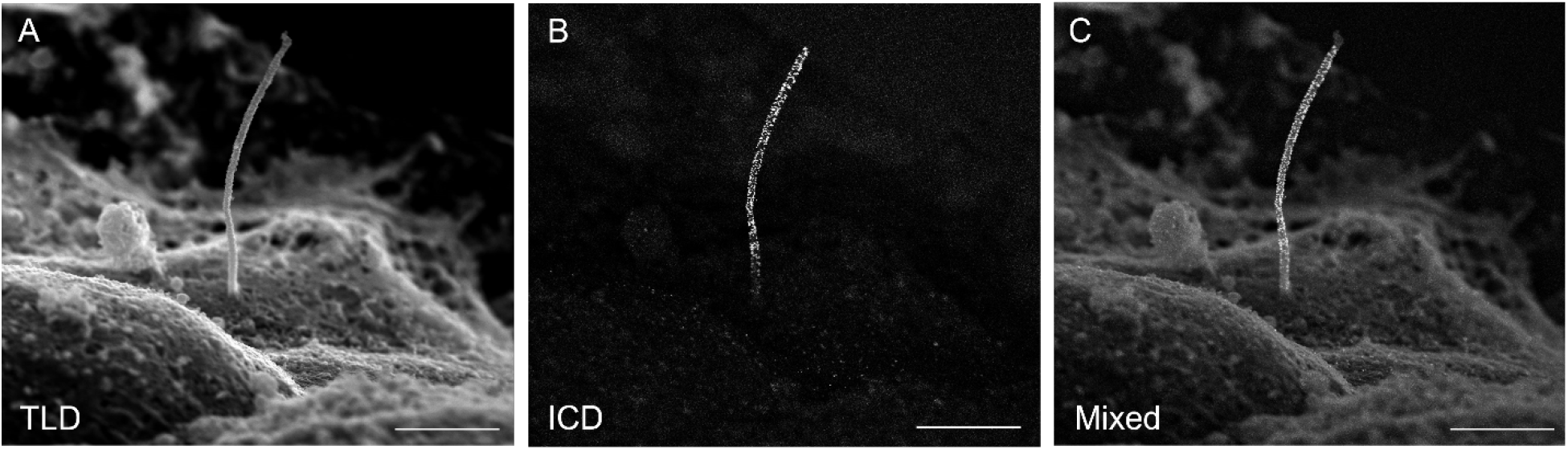
Multi-detector scanning. Simultaneous capture of secondary and backscatter electron images of a single mouse islet primary cilium labeled with acetylated alpha-tubulin tubulin (Proteintech 66200-1-Ig, mouse IgG1, 1:100). (**a**) Secondary electron imaging with Through Lens Detector (TLD), (**b**) backscatter electron imaging using In Column Detector (ICD), (**c**) combined images are generated through digital mixing of TLD and ICD images rather than through a specific detector. In general, mixed images provides the best visualization of immunogold position against islet surface textures and are used for most figures in this report. Scale, 2 μm.

**Figure 3.**
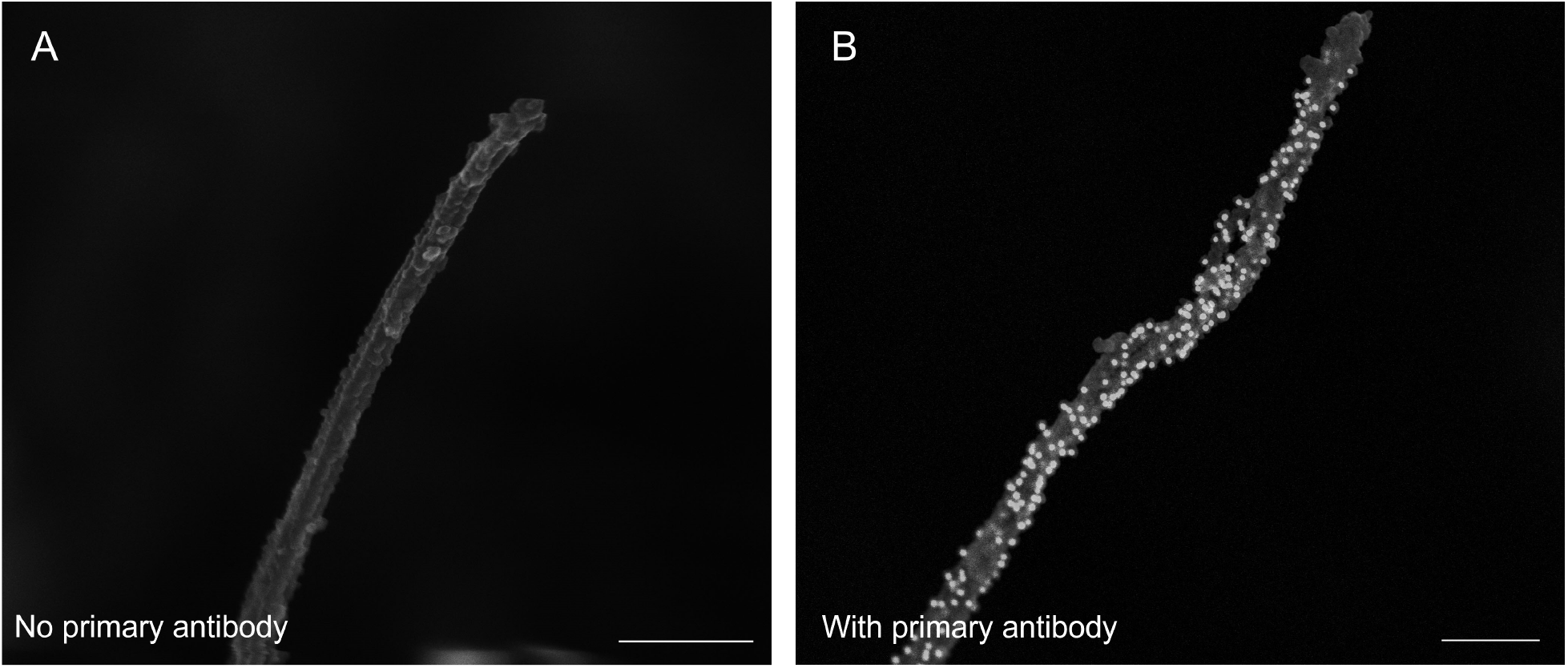
Negative control for immuno-SEM. Human islet cilia labeled without (**a**) or with (**b**) primary antibody to acetylated alpha-tubulin (Proteintech 66200-1-Ig, mouse IgG1, 1:100). The no-primary antibody condition controls for nonspecific binding of the secondary gold antibody (Jackson ImmunoResearch, donkey-anti-mouse #715-215-150, 1:20). Scale 400 nm.

Antibody performance for immuno-SEM was comparable to their performance in IF imaging, with core axonemal tubulin being most abundantly detectable. Acetylated alpha tubulin is a universal marker of primary cilia across tissue types and in our study reliably labeled the entire ciliary axoneme in mouse and human islets (**Figure 2-4**). A second post-translational modification of tubulin, polyglutamylated tubulin, was also readily detected through the entire length of the cilium (**Figure 5**). Intraflagellar transport IFT88 was detected in a scattered pattern along the entire axoneme, using an affinity-purified rabbit polyclonal antibody previously validated by IF^13^ in islet cilia (**Figure 6**). We did not observe discrete IFT train-like structures corresponding to IFT88 immunolabeling on the axonemal surface and reason that the train complexes may have been partially disrupted by the demembranation process.

**Figure 4.**
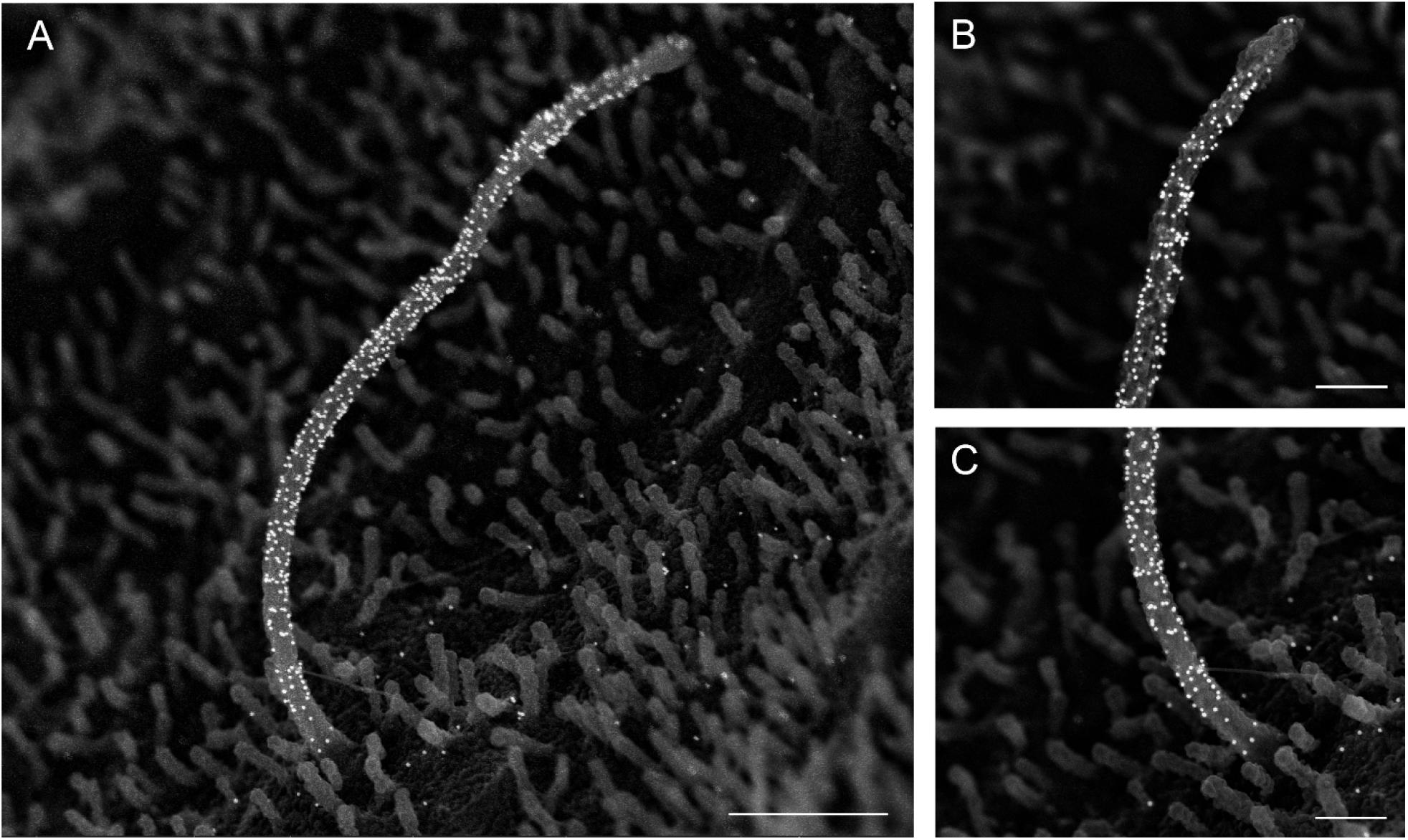
Acetylated alpha tubulin labeling in human islet cilia. Human islet primary cilium labeled with mouse-anti-acetylated alpha tubulin (Proteintech 66200-1-Ig, mouse IgG1, 1:100), showing clear and abundant labeling on the ciliary axoneme, in contrast to low background labeling on the cell surface. Images with mixed secondary and backscattered electron signal. (**a**) whole cilium, scale 1 μm, (**b**) cilia tip, scale 400 nm, (**c**) cilia base, scale 400 nm.

**Figure 5.**
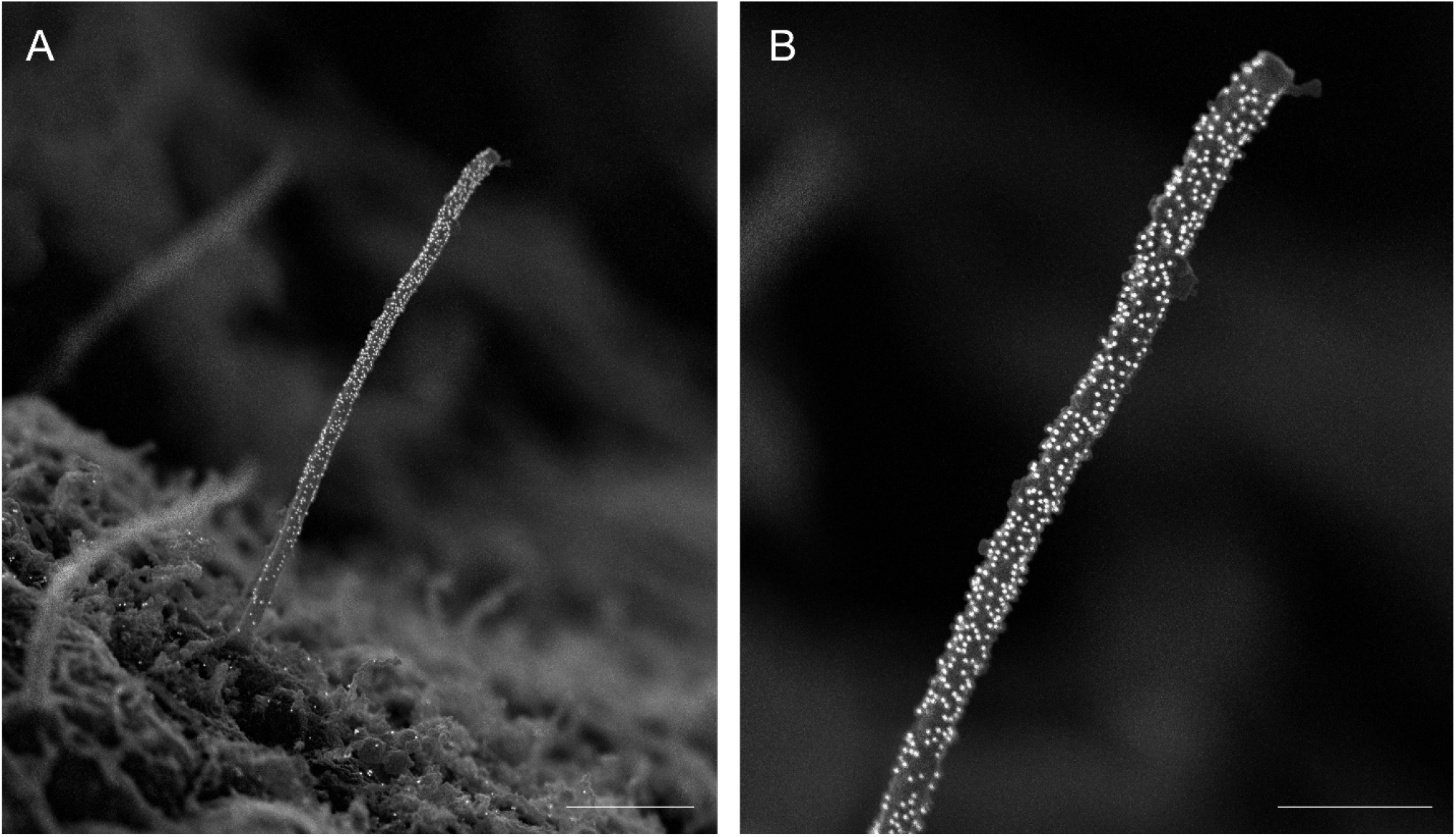
Polyglutamylated tubulin labeling in mouse islet cilia. Another post-translational modification of the ciliary axoneme as labeled by mouse-anti-polyglutamylated tubulin (GT335 antibody, AdipoGen, mouse IgG1k 1:400). (**a**) whole cilium, scale 1 μm, (**b**) distal cilium and tip, scale 500 nm.

**Figure 6.**
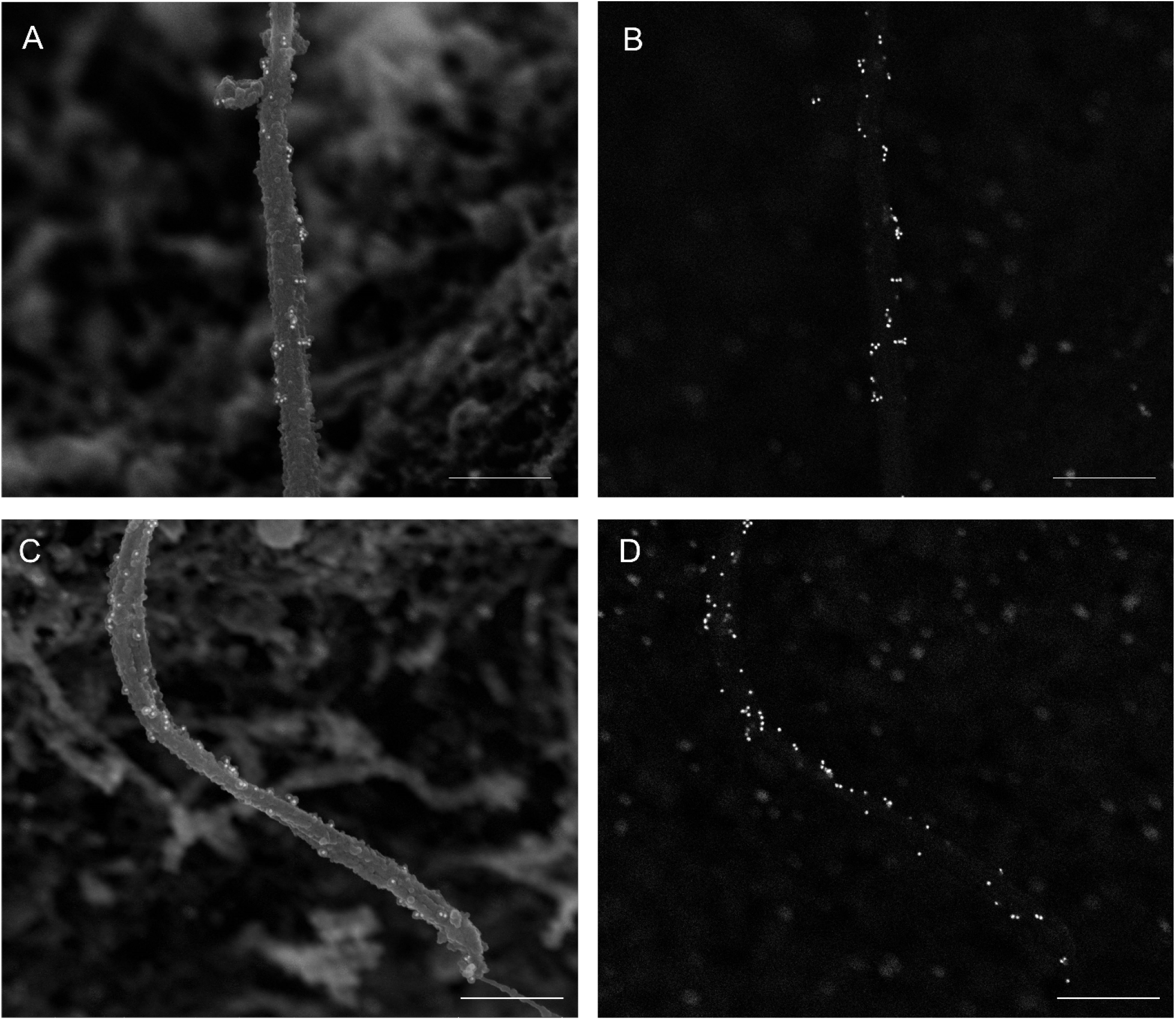
Intraflagellar transport 88 (IFT88) in mouse islet cilia. Mix images (**a, c**) and corresponding ICD images showing backscatter electron signal (**b, d**) of IFT88 labeling in mouse islets (rabbit-anti-IFT88, NSJ Bio F41236, 1:50). Scale 500 nm.

We did observe variable performance among antibodies for the same antigen, even those validated across other immunolabeling protocols. Arl13b is a small ciliary G protein of the Ras superfamily and is a standard ciliary protein marker for IF studies. We tested two commercial Arl13b antibodies that our lab relies on for IF imaging. One antibody (Neuromab N295B/66, 1:100 dilution) gave clear ciliary labeling with near zero background signal on the membrane (**Figure 7**), while another antibody (Proteintech 17711-1-AP, 1:100 dilution) despite being a strong performer for IF gave such heavy background signal in immuno-SEM that rendered its ciliary labeling uninterpretable (**Figure 8**). This was a surprise given clean IF staining by the Proteintech antibody in our experience and highlights the need to empirically test each antibody for immuno-SEM despite prior validation by other imaging modalities.

**Figure 7.**
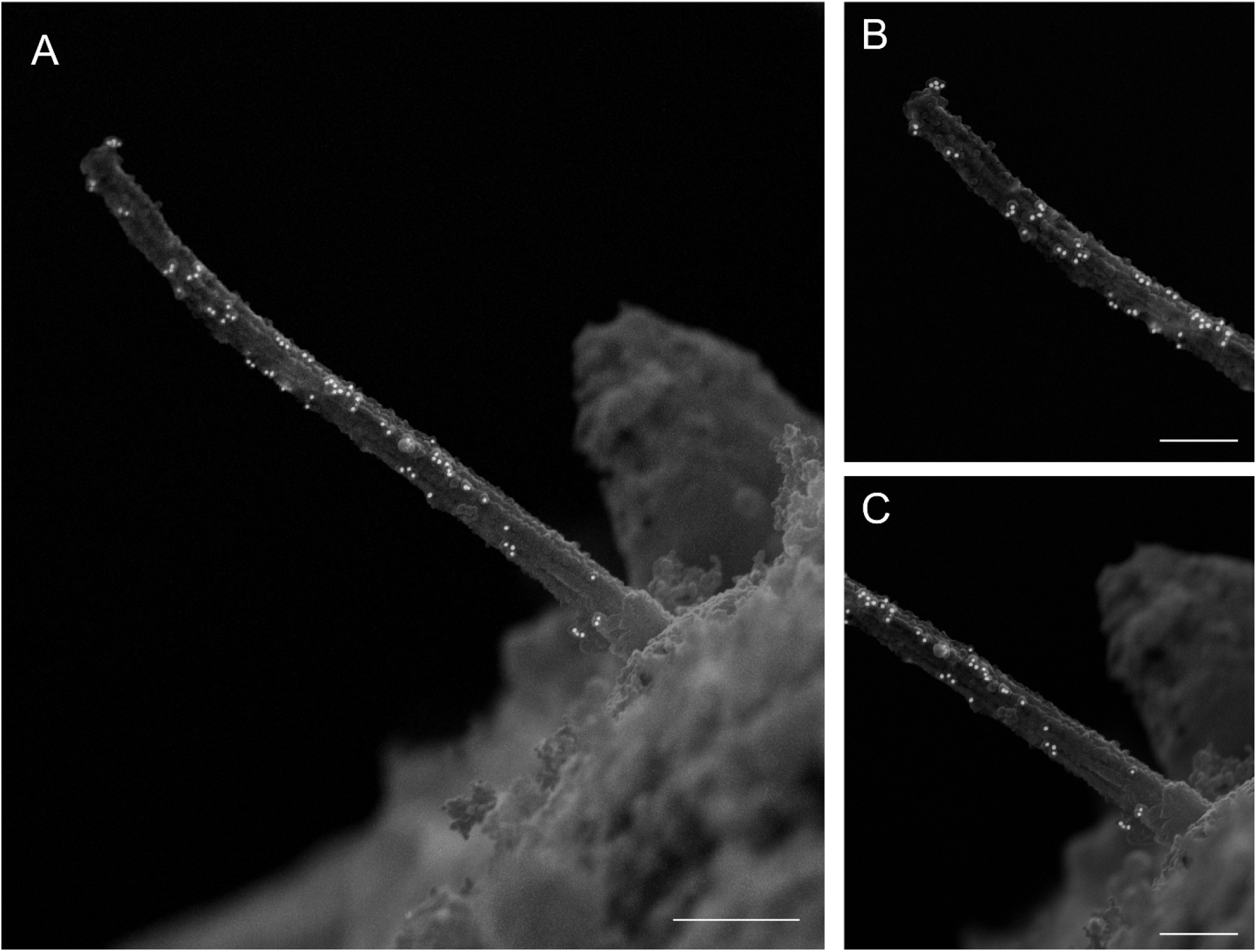
Arl13b labeling in mouse islet cilia. Clean Arl13b immunolabeling is achieved using a monoclonal primary antibody (NeuroMab N295B/66, mouse IgG2a, 1:100). (**a**) Whole cilium, scale 1 μm, (**b**) distal cilium and tip, scale 500 nm.

**Figure 8.**
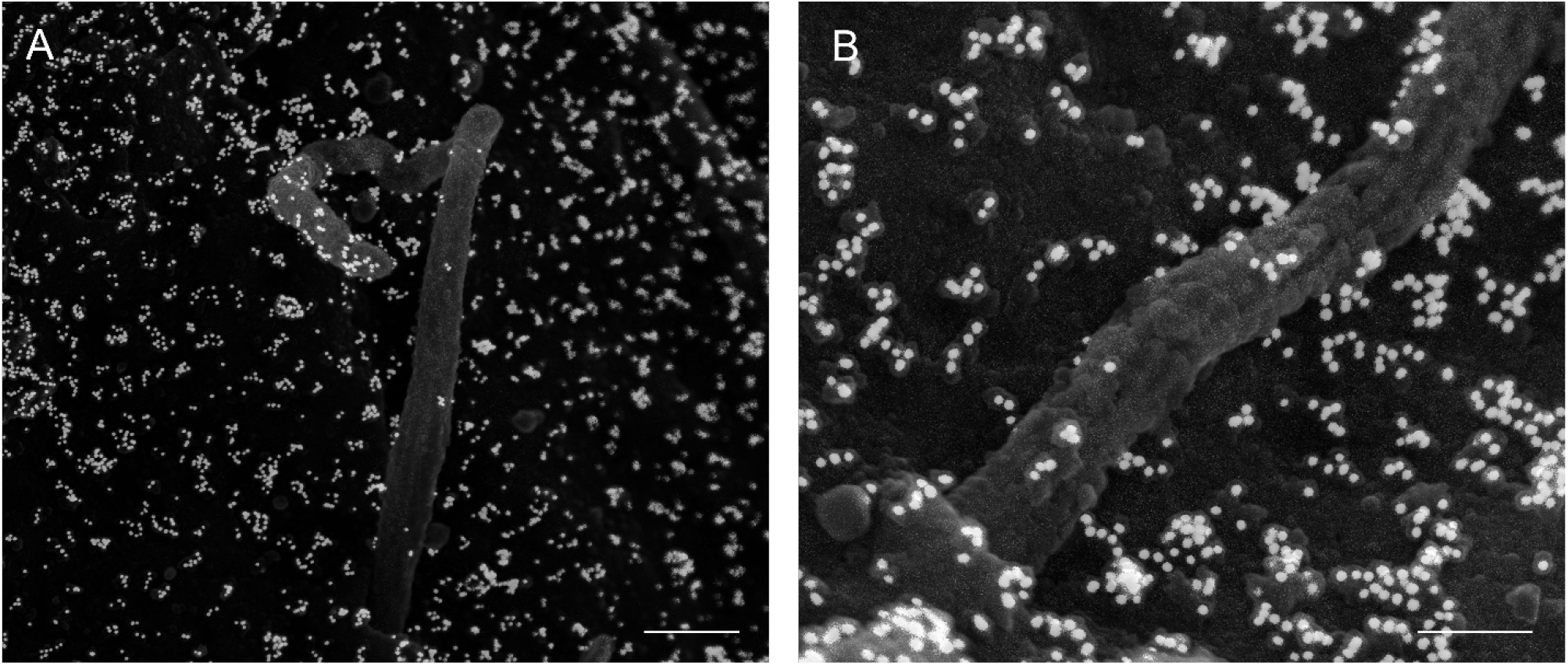
Arl13b labeling, alternate antibody with high background. A rabbit polyclonal Arl13b antibody (Proteintech 17711-1-AP, 1:100) produced heavy background labeling on the mouse islet surface, making it difficult to interpret ciliary signals. (**a**) Whole cilium, scale 500 nm, (**b**) ciliary base, scale 200 nm.

As we and others have observed motile properties in primary cilia^5,7,8^, we tested the presence of axonemal dynein in islets. Two axonemal dynein antibodies, dynein heavy chain 5 (DNAH5) and dynein intermediate chain 1 (DNAI1), produced specific labeling on the ciliary axoneme with clean background (**Figure 9**). These immuno-SEM localizations in mouse islets, together with our prior IF data showing axonemal dynein expression in islet cilia using these same antibodies in human samples^8^, demonstrate the presence of motor dynein in islet primary cilia and support the notion that cilia motility may be a conserved feature across species.

**Figure 9.**
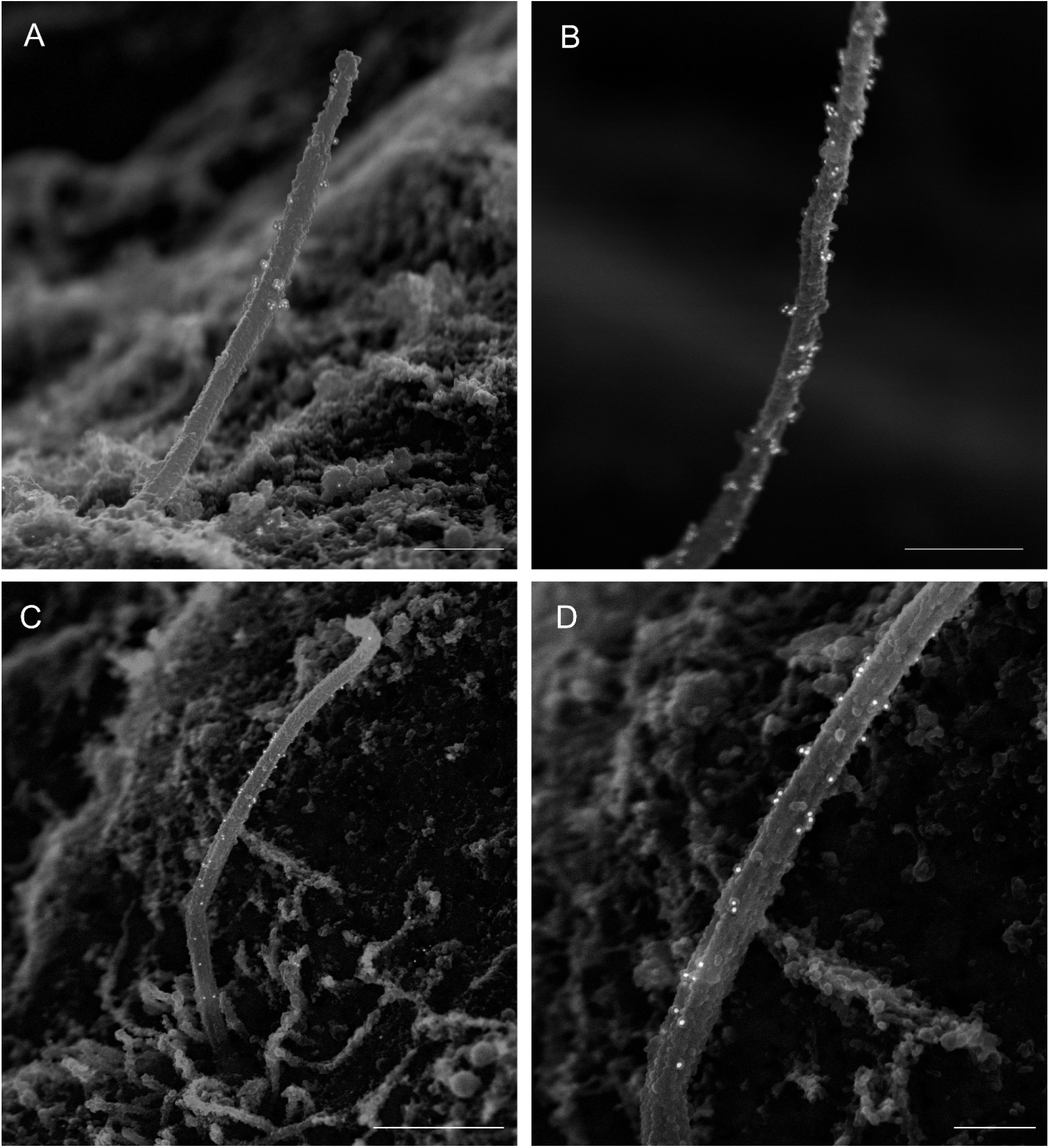
Axonemal dynein in mouse islet cilia. Presence of motor dynein in the primary cilia axoneme as indicated by immunogold labeling of dynein heavy chain 5 (rabbit-anti-DNAH5, Sigma HPA 037470, 1:100) and dynein intermediate chain 1 (mouse-anti-DNAI1, NeuroMab 75-372, 1:100). (**a**) DNAH5, whole cilium, scale 500 nm, (**a**) DNAH5, distal cilium, scale 500 nm, (**c**) DNAI1, whole cilium, scale 1 μm, (**d**) DNAI1, mid-axoneme, scale 400 nm.

## DISCUSSION

Our study demonstrates that the immuno-SEM technique is suitable for protein identification in islet primary cilia using standard commercial antibodies. Whereas compelling protein localization data is often unattainable by immunofluorescence (IF) for reasons discussed in the introduction, we found immuno-SEM to offer the unique capability to demonstrate ciliary protein distribution while resolving ultrastructural details. A key advance in our protocol is that, with demembranation, our method can be applied to labeling core axonemal proteins and not limited to proteins on the ciliary surface. We discuss below a number of technical details key to the success and reproducibility of our immuno-SEM workflow.

### Sample stability and preservation

Sample detachment is a common problem for primary tissues such as islets, leading to attrition of imageable specimens throughout the workflow. This led to increased time required for sample prep and, often, for experiments to be repeated. In our experience, islets that were loosely attached were also unusable, as these had poor grounding and conductivity and therefore led to charging issues during imaging. For best success, we found that coating the glass coverslips with recombinant human laminin (Gibco rhLaminin-521) and allowing the islets to fully adhere, up to 3 days, helped ensure that adequate islet numbers are preserved. Islet attachment and quality is routinely assessed throughout the workflow to ensure proper concentration and distribution on the grid.

### Fixation

Immuno-SEM is a hybrid protocol between immunohistochemistry and scanning EM, therefore requires multiple chemical fixation steps dedicated for these different procedure segments. The choice of fixatives is important when considering antibody performance, just as it has been demonstrated for cilia IF imaging^16^. The requisite fixative for EM is glutaraldehyde (GA), which in high concentrations is incompatible with antibody labeling yet must be included upfront to preserve ultrastructural morphology and at the end to crosslink primary-secondary antibody interaction. We found that islets could tolerate 0.25% glutaraldehyde (GA) during the demembranation step prior to antibody staining. We then rinse and fix the samples with 4% PFA as would for routine immunohistochemistry, and after antibody staining is complete, do a final fix with 2% GA for electron microscopy.

### Antibody performance

Antibody performance is essential to the success of immuno-SEM and depends on critical factors such as sample quality, antigen concentration and availability, fixative choice, and labeling conditions. Generally, strong performance by an antibody in IF is a good indication that it may work for immuno-SEM. Still, this requires a certain amount of trial and error, as can be seen in our study, and optimization is needed to identify successful antibody dilution and staining conditions^17^. We found a number of steps to enhance primary antibody labeling: 1) glycine quenching, which increases signal specificity by reducing non-specific free aldehyde binding to the antibody^10^, 2) membrane-stripping, which exposes antigens below the ciliary surface, and 3) use sufficient antibody concentrations, usually 5-10x higher than for IF, which is required to survive extensive washing and multiple fixatives. For secondary antibodies, we used commercial 12 nm colloidal gold-conjugated Ig to mouse and rabbit (Jackson ImmunoResearch) and found these to produce robust and reliable labeling.

### Inclusion of controls

The inclusion of proper negative and positive controls is essential to the rigor of any imaging study, immuno-SEM included. Unlabeled samples such as no-primary antibody provide good controls against nonspecific labeling by secondary colloidal gold. Of note, this does not demonstrate specificity of primary antibody binding, which would ideally be done by pre-incubating the antibody with the antigen peptide, replacing the primary antibody with a nonimmune Ig of the same isotype, or by validating in gene knockout samples. For positive control, the use of alternative targeted antibodies to the same antigen and especially from a different host species would be a good way to validate labeling results.

### Continuity of workflow

While the sample adhering and immunolabeling steps offer flexibility on timing and duration, certain parts of the protocol should be performed without interruption, otherwise could lead to processing issues. We found that mouse islets were susceptible to damage if they had a longer pause between critical point drying (CPD) and coating, which would lead to severe charging issues and make cilia difficult to image in high-resolution mode. This likely stems from moisture accumulation and trapping under the coating, which could be minimized by coating samples immediately after CPD and storing them in a desiccator until imaging. Human islets typically behaved more stably and did not have as many processing issues, but had much fewer cilia on their surface, thus also required care during the workflow to not reduce sampling number.

### Sample charging and coating

Electric charging is a common problem during imaging, an issue stemming from the production of excess secondary electrons resulting in the sample surface becoming positively charged. The plasma of secondary electrons that forms on the surface of the sample interferes with further interaction of the incident beam with the sample which results in charging effect and inability to keep electron beam in focus. In our islet samples, we frequently needed to image multiple areas of the grid because of charge built-up in certain regions. To combat charging, we used three strategies: 1) low levels of osmium in the sample prep, 0.5% instead of the usual 2% used for TEM/SEM – the latter would make samples better grounded but given the high atomic number of osmium would generate excess backscatter signal to interfere with the gold label; 2) carbon coating, which deposits a thin layer of conductive material to reduce charge build-up by grounding excess electrons. We used 12 nm carbon rather than heavier metals such as platinum and chromium^18^ which would have generated too much backscatter; 3) limiting the accelerating voltage during imaging, e.g. 5 kilovolts (kV), which is on the high end for biological samples but was the level necessary on our Helios FIB-SEM system for producing sufficient backscatter electron signal to see the gold. Some of these technical adjustments led to small changes to the specimen itself, for example carbon coating adds 12 nm to the sample surface which slightly increases their physical dimensions, which should be accounted for when extrapolating size information. Carbon coating also likely contributed to “gumming up” of the sample surface and limited our ability to resolve axonemal details and visualize gaps between microtubules, a caveat to be kept in mind for image interpretation.

In summary, we show feasibility of the immuno-SEM technique for identifying and visualizing primary cilia proteins on pancreatic islets. This workflow performs robustly for multiple protein antigens tested in our study and across two species, mouse and human. We emphasize the need to optimize labeling conditions in each antibody, cell, and tissue type, rather than strict adherence to our protocol. We hope that discussion of the critical parameters in sample preparation and imaging will facilitate future immuno-SEM studies to enable protein characterization of primary cilia.

## ACKNOWLEDGEMENT

Sanja Sviben is currently a Scientist at the Stanford University Cryo-Electron Microscopy Center, Stanford University, Palo Alto, CA. Alexander Polino is currently a Scientist in Oncology at Avantor in Philadelphia, PA. We gratefully acknowledge funding for this study including NIH grants DK115795, DK127748, and DK138974 to J.W.H. The Washington University Center for Cellular Imaging (WUCCI) is supported by Washington University School of Medicine, The Children’s Discovery Institute of Washington University and St. Louis Children’s Hospital (CDI-CORE-2015-505 and CDI-CORE-2019-813), the Foundation for Barnes-Jewish Hospital (3770 and 4642) and the Washington University Diabetes Research Center (DK020579). Human pancreatic islets from the Integrated Islet Distribution Program (IIDP) (RRID:SCR_014387) were supported by NIH grant #2UC4DK098085 and the JDRF-funded IIDP Islet Award Initiative.

## Author Contributions

S.S. and J.W.H. conceptualized the study. J.W.H.’s laboratory provided islet specimens. A.J.P., I.M., and S.S. performed sample preparation. S.S. performed scanning electron microscopy. J.W.H. wrote and revised the manuscript with input from all authors.

## Competing interests

The authors declare no competing interests.

## Notes

### Competing Interest Statement

The authors have declared no competing interest.

